# End Binding protein 1 promotes specific motor-cargo association in the cell body prior to axonal delivery of Dense Core Vesicles

**DOI:** 10.1101/2023.01.12.523768

**Authors:** Junhyun Park, Kenneth G. Miller, Pietro De Camilli, Shaul Yogev

## Abstract

Axonal transport is key to neuronal function. Efficient transport requires specific motor-cargo association in the soma, yet the mechanisms regulating this early step remain poorly understood. We found that EBP-1, the *C. elegans* ortholog of the canonical microtubule end binding protein EB1, promotes the specific association between kinesin-3/KIF1A/UNC-104 and Dense Core Vesicles (DCVs) prior to their axonal delivery. Using single-neuron, *in vivo* labelling of endogenous cargo and EBs, we observed reduced axonal abundance and reduced secretion of DCV cargo, but not other KIF1A/UNC-104 cargo, in *ebp-1* mutants. This reduction could be traced back to fewer exit events from the cell body, where EBP-1 colocalized with the DCV sorting machinery at the trans Golgi, suggesting that this is the site of EBP-1 function. In addition to its microtubule binding CH domain, mammalian EB1 interacted with mammalian KIF1A in an EBH domain dependent manner, and expression of mammalian EB1 or the EBH domain was sufficient to rescue DCV transport in *ebp-1* mutants. Our results suggest a model in which kinesin-3 binding and microtubule binding by EBP-1 cooperate to transiently enrich the motor near sites of DCV biogenesis to promote motor-cargo association. In support of this model, tethering either EBP-1 or a kinesin-3 KIF1A/UNC-104 interacting domain from an unrelated protein to the Golgi restored the axonal abundance of DCV proteins in *ebp-1* mutants. These results uncover an unexpected role for a microtubule associated protein and provide insight into how specific kinesin-3 cargo are delivered to the axon.

## Introduction

Axonal transport necessitates cargo-motor association and efficient cell body exit prior to long-range runs, but these early steps of transport remain poorly understood. The major axonal motor kinesin-3/KIF1A/UNC-104 is thought to be activated in the cell body following relief of autoinhibition and interaction of its PH domain with phospholipids on cargo vesicles.^1–5^ However, UNC-104/KIF1A can transport different cargo types, raising the question of whether additional mechanisms exist that could promote the transport of specific cargo.^6–11^

A case in point is synaptic vesicle precursors (SVPs) and dense core vesicles (DCVs), both UNC-104/KIF1A cargo, which differ from each other in terms of membrane composition and the dependence on different biogenesis and sorting mechanisms in the cell body.^12–19^ Interestingly, recent work in *C. elegans* showed that the cargo adaptor complexes which sort DCVs and SVPs for axonal delivery (UNC-101/AP-1μ and APM-3/AP-3μ, respectively) are found on separate sub-domains of the trans Golgi.^14, 20^ This finding raises the possibility that the association of UNC-104/KIF1A with specific cargo could be regulated by proteins that reside at these spatially segregated biogenesis sites.

End Binding (EB) proteins are a highly conserved family of microtubule (MT) associated proteins (MAPs) that bind the growing MT end.^21^ EBs are characterized by an N-terminal calponin homology (CH) domain, which mediates MT end tracking, and a C terminal End-binding Homology (EBH) domain, which mediates interactions with EB binding proteins.^22–25^ EBs can recruit MTs to membranous organelles and in neurons they are thought to play important roles in controlling microtubule organization, either by regulating microtubule dynamics, mediating binding to structures such as the axon initial segment (AIS) or guiding MT growth along other microtubules or the cortex.^26–29^ EBs also play a role in the initiation of retrograde transport from the tip of the axon by facilitating dynein recruitment and may compete with kinesin-3 for the GTP cap at presynapses.^30–32^ Unexpectedly, recent work revealed that cultured mammalian cells in which all three EBs were disrupted show relatively mild mitotic defects.^29, 33^ Similarly, worm embryos lacking all EBs are viable and develop normally.^34^ Hence, despite the extensive characterization of EBs at the structural, biochemical and cell biological levels, the full range of their functions *in vivo* remains to be elucidated.

Here, using single-neuron, endogenous labelling of cargo and EBs *in vivo*, we found a role for the *C. elegans* EBP-1 in promoting KIF1A/UNC-104 dependent axonal delivery of DCVs but not SVPs or ATG-9 vesicles. EBP-1 is enriched near the trans Golgi with the DCV sorting machinery and *ebp-1* mutants showed reduced numbers of DCVs exiting the cell body, suggesting that this is its relevant site of action. Consistently, DCV abundance in the axons of *ebp-1* mutants could be restored by expression of a Golgi tethered fragment of EBP-1. We find that mammalian EB1 could rescue *ebp-1* mutants – suggesting conservation of its function – and its EBH domain co-immunoprecipitated with kinesin-3/KIF1A. Loss and gain of function experiments indicate that EB promotes DCV transport through the microtubule binding CH domain and the UNC-104/KIF1A binding EBH domain, with the latter playing a more prominent role. We propose through binding UNC-104, and potentially through the ability of the CH domain to interact with MT ends, EBP-1 transiently enriches the motor near DCV sorting sites, thus promoting specific motor-cargo association and efficient transport. In support of this model, EBP-1 function could be replaced by a Golgi-tethered UNC-104/KIF1A interacting domain from an unrelated protein, SYD-2.

## Results

### Loss of EBP-1 results in a specific loss of Dense Core Vesicle cargo in the axon

Recent *in vitro* studies highlighted the role of MAPs, including EBs, as potential regulators of molecular motors.^30^ To investigate the role of End-Binding proteins in neuronal transport *in vivo*, we used the *C. elegans* bipolar cholinergic motor neuron DA9. The cell body of DA9 resides in the preanal ganglion, from where the axon grows posteriorly, traverses to the dorsal cord and grows anteriorly to the mid-body. Following an asynaptic region of ~45 μm in the dorsal cord, *en passant* presynaptic boutons form onto dorsal muscle at a stereotypic stretch of 70~90 μm (**Figure 1A**).^35, 36^ A DCV luminal cargo, INS-22∷Emerald, expressed under control of a DA9 promoter, shows enrichment at the synaptic region and is also detected in asynaptic regions, consistent with previous reports (**Figure 1C**).^37, 38^

**Figure 1:**
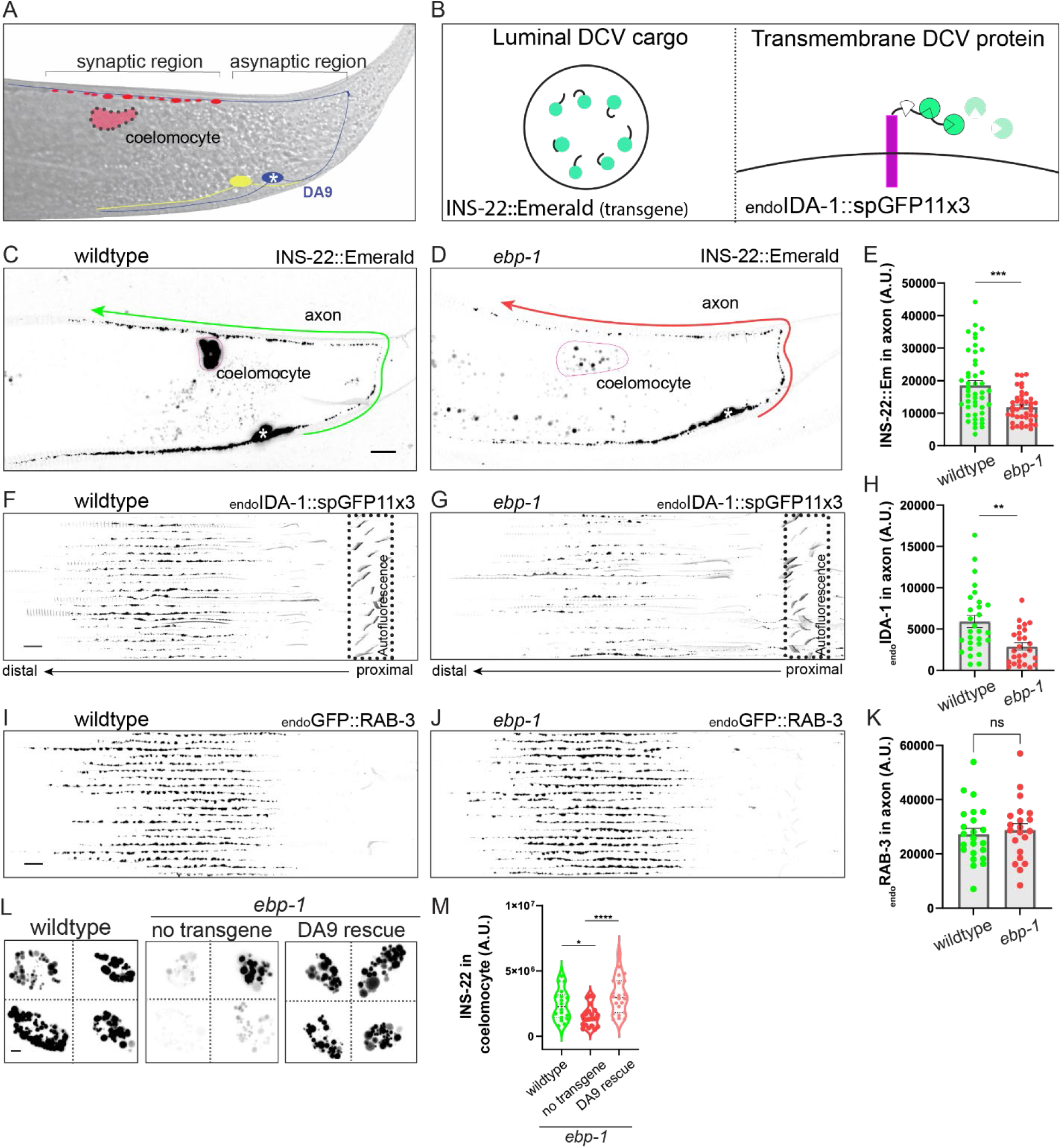
Loss of *ebp-1* results in a specific loss of dense core vesicles from the axon. (A) Schematic of the DA9 motor neuron. DA9 axon traverses dorsally and forms en passant presynapses (red boutons). White asterisk indicates the soma in all figures. The ccDL coelomocyte, adjacent to DA9 synapses is in pink. VA12 neuron (yellow) synapses on the ventral side and is also labeled by *Pmig-13* promoter. (B) Diagram of INS-22, luminal DCV cargo, transgene (left box) and splitGFP tagging strategy for labeling endogenous transmembrane DCV cargo, IDA-1 (endoIDA-1, right box). (C,D) Confocal images of wildtype (C) and *ebp-1* mutant (D) expressing INS-22∷Emerald. Green and red arrows points toward the more distal axon of DA9 and the straightened axon that spans the arrow line was quantified. Pink silhouette outlines the coelomocyte. Scale bar 10 μm. (E) Quantification of (C) and (D). (F-H) Twenty axons per genotype were straightened and aligned from cell body (proximal) to past the synaptic region (distal). Signal is from splitGFP tagged to endogenous IDA-1. Non-specific background fluorescence from the rectum (diagonal lines) is indicated by dotted box (F,G). Quantification of fluorescence in (H). Scale bar 10 μm. (I-K) Fluorescence of endogenously labeled RAB-3 vesicles in wildtype and *ebp-1* mutant (I, J), quantified in (K). Scale bar 10 μm. (L-M) Representative images of INS-22∷Emerald in coelomocyte in wildtype, *ebp-1* mutant, and DA9 specific expression of EBP-1 in *ebp-1* mutant. Scale bar 1 μm. Quantification of fluorescence in (M) n=21-46. ****p<0.0001; ***p<0.001; **p<0.01, *p<0.05 (t-test for (E), (H), (K) and one-way ANOVA for (M)).

Examination of a previously described deletion allele of *ebp-1* (*he279)* or an allele generated for this study (*wy1156*), revealed a ~50% reduction of INS-22∷Emerald in the axon of DA9 (**Figure 1D** and **1E**), suggesting that EBP-1 is required for the normal abundance of DCVs in the axon (note that the deletions in the *wy1156* and *he279* alleles encompass both *ebp-1* and the neighboring gene, *ebp-3*, which is likely a pseudogene.^34^ We therefore refer to *ebp-1, ebp-3* mutants as *ebp-1*. All alleles used are null unless indicated otherwise; deletions of specific domains are noted by Δ).

To determine whether other DCV cargo requires EBP-1, we used CRISPR to tag the C terminus of IDA-1/phogrin, a known transmembrane DCV protein, at its endogenous locus with spGFP11 (referred to as endoIDA-1) and visualized it in DA9 at native levels by expressing spGFP1-10 under *Pmig-13* promoter (**Figure 1B**). endoIDA-1 was more enriched at synapses compared to INS-22 in wildtype animals and showed a similar ~50% reduction in *ebp-1* mutants (**Figure 1F**, **1G**, and **1H**). These results indicate that both luminal and transmembrane DCV cargo require EBP-1 for their axonal localization.

Given that the DCVs are prominently lost in *ebp-1* mutants from the dorsal axon where presynaptic sites lie, we tested whether their loss may be due to excessive secretion. We generated double mutants between *ebp-1* and *unc-31/CAPS*, which is required for DCV secretion.^37, 39, 40^ If *ebp-1* mutants lead to excessive secretion, we would expect *unc-31* mutants to restore DCVs to the axon. However, *ebp-1; unc-31* double mutants showed ~40% loss of the INS-22 signal compared to *unc-31* mutants (**Figure S1C** and **S1D**), suggesting that loss of DCV cargo from the DA9 axon of *ebp-1* mutants is caused by reduced delivery of DCVs, not over secretion of their contents.

To examine how reduced axonal DCV abundance in *ebp-1* mutants affects DCV cargo secretion, we measured INS-22 levels in coelomocytes, which are scavenger cells that endocytose the secreted INS-22 (Pink outline; **Figure 1A** and **1C**). Measuring the fluorescence of INS-22 endocytosed by these cells serves as a proxy of secreted INS-22 over time and has been a replicable metric for DCV secretion in *C. elegans*.^41^ Quantification of INS-22∷Emerald fluorescence in the closest coelomocyte to DA9 synapses revealed a 50% reduction in *ebp-1* mutants compared to wildtype (**Figure 1L**). This effect could be rescued by cell specific expression of EBP-1 in DA9 (**Figure 1L** and **1M**), indicating that the reduced INS-22 signal in coelomocytes is due to a reduction in INS-22 secreted from DA9 and not to a defect in coelomocyte function. DCV cargos play important roles in the animal physiology. In *C. elegans*, neuropeptides both promote and inhibit egg-laying; specifically, ectopic expression of the neuropeptide NLP-3 was shown to induce premature egg-laying, leading to the retention of very few eggs in the animal’s uterus.^42, 43^ Consistent with a role for EBP-1 in promoting DCV delivery for secretion, we found that *ebp-1* mutants suppressed the excessive egg-laying behavior of animals overexpressing NLP-3 (**Figure S1A** and **S1B**). Collectively, these results indicate that loss of *ebp-1* leads to reduced axonal abundance of DCVs and reduced secretion of DCV cargo.

DCVs are transported to the axon by kinesin-3/KIF1A/UNC-104, which also transports synaptic vesicle precursors (SVPs) and other synaptic cargo to the presynapses.^6–8, 10, 11^ To determine if the DCV phenotype of *ebp-1* mutants reflects a general reduction in UNC-104 activity, we measured the axonal signal of RAB-3, a SVP marker, labelled at its endogenous locus with a FLP-ON GFP cassette and visualized with a DA9 FLP. Surprisingly, the endoRAB-3 signal was not decreased in *ebp-1* mutants (**Figure 1I**, **1J**, and **1K**). The levels of other UNC-104 cargo, the transmembrane SVP protein SNG-1/synaptogyrin and the autophagic regulator ATG-9 were also unaffected in *ebp-1* mutants (**Figure S1E**, **S1F**, **S1G**, and **S1H**). These results suggest that *ebp-1* is specifically required for DCV trafficking.

We note that in *ebp-1* mutants, the spacing between adjacent presynaptic boutons was reduced, a phenotype which could be best visualized with the active zone marker CLA-1 (**Figure S2H** and**S2I**).^45^ However, as described below in structure function analysis and rescue experiments, this phenotype stems from a separate function of EBP-1 that is not related to DCV delivery and is not pursued in this study.

### The EBH domain is sufficient to restore axonal DCVs in *ebp-1* mutants

To gain further insight into how EBP-1 functions in DCV trafficking, we assayed the requirement for different EB proteins and domains in DA9. We first examined the EBP-1 paralog, EBP-2, which is expressed in DA9 at similar levels to EBP-1 (**Figure S2F**). To assess the DCV loss phenotype, we measured overall fluorescence in the axon as well as the number of IDA-1 puncta in the synaptic region. *ebp-2* (*he278*) deletion mutants showed only a minor reduction in axonal endoIDA-1 compared to wildtype, but that reduction was not on par with the phenotype observed in *ebp-1* mutants (**Figure S2A, S2B**. and **S2C**). Furthermore, *ebp-2* mutants did not enhance the phenotype of *ebp-1* mutants (**Figure S2B**, **S2C, S2D**, and **S2E**). These results suggest that EBP-2 plays a relatively minor role in DCV trafficking.

Next, we used CRISPR to delete either the CH domain (*ebp-1ΔCH*) or the C terminus which contains the EBH domain and acidic tail (*ebp-1ΔC*; **Figure 2A**) from the *ebp-1* genomic locus. In parallel, we tagged the resulting EBP-1 fragment at their endogenous loci with spGFP11 and confirmed that they were still expressed in DA9 with *Pmig-13* driving spGFP1-10 (**Figure S2F**). Interestingly, both deletion mutants did not show a significant loss of endoIDA-1 from the axon (**Figure 2C** and **2D**). In contrast, *ebp-1ΔCH* did lead to a reduced distance between synaptic boutons but *ebp-1ΔC* did not (**Figure S2H and S2I**), indicating that this phenotype is separate from the loss of axonal DCVs. Many EB binding proteins associate with EBs via a SxIP motif, which binds the EBH domain.^23, 29^ However, two point-mutations that are predicted to eliminate binding to SxIP motif proteins (Y264A and E272A) did not alter axonal DCV levels (**Figure 2F** and **2G**). ^23, 24^ In contrast, deleting both the CH domain and the C-terminus phenocopied *ebp-1* deletion (**Figure 2E**). A CH deletion with Y264A and E272A also phenocopied *ebp-1*, but the resulting protein was expressed at significantly lower levels (**Figure 2F, 2G**, and **S2F**), preventing us from drawing definitive conclusions about Y264A and E272A in DCV transport. Together, these results suggest that both MT binding and an association with an EBP-1 binding protein need to be eliminated to mimic *ebp-1* deletion mutants. One possible interpretation of this result is that the relevant EBP-1 binding protein is also able to bind MTs, as seen with several EBP-1 interactors.^46–48^

**Figure 2:**
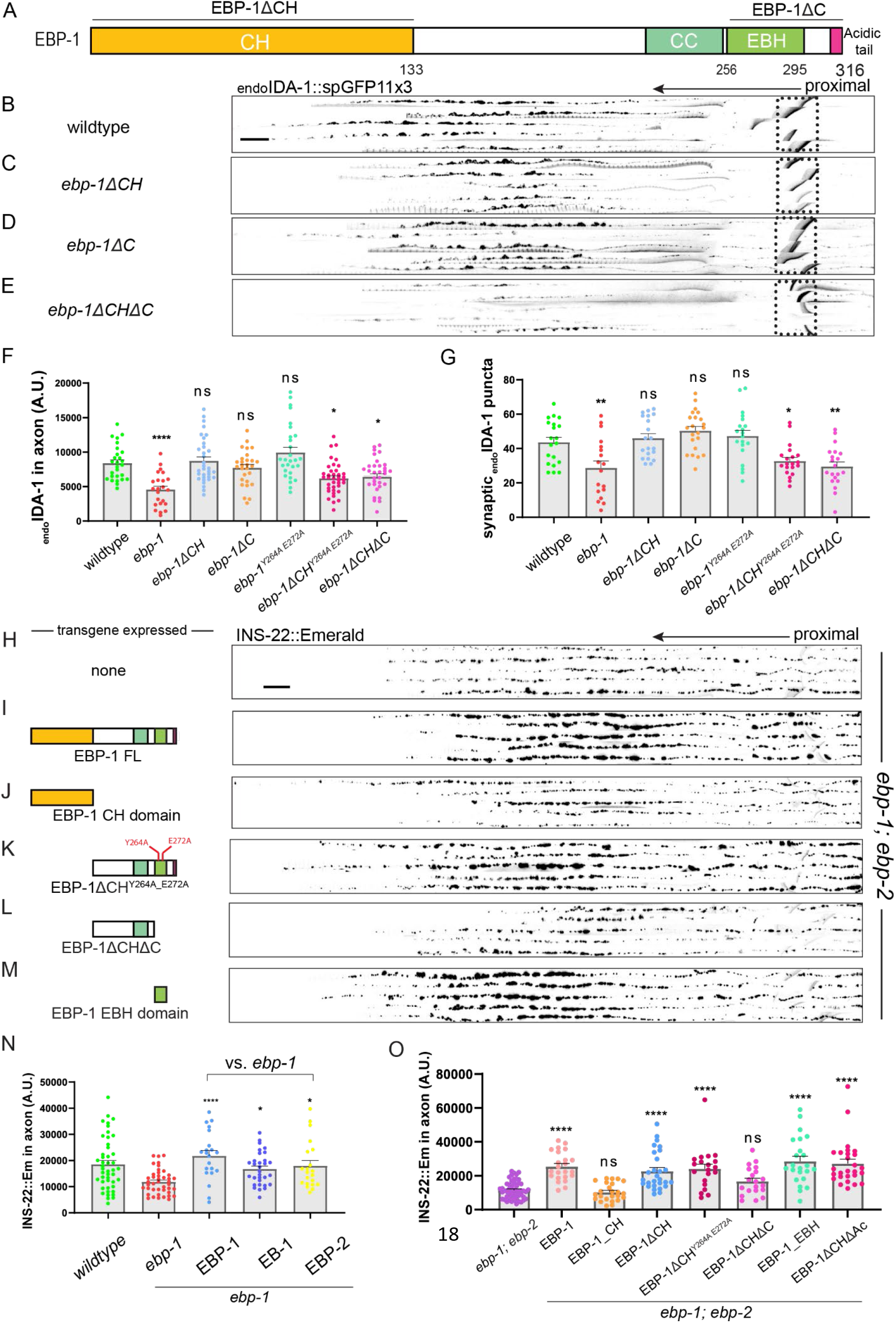
The EBH domain is sufficient for proper DCV trafficking. (A) Diagram of EBP-1 and its predicted domains. EBP-1ΔC truncates both EBH domain (lime green) and acidic tail (magenta). (B-E) Alignments of 5 representative axons per genotype showing endoIDA-1∷GFP in DA9. Black arrow points distally. Autofluorescence from rectum in dotted box. Scale bar 10 μm. (F) Quantification of fluorescence of endoIDA-1. (G) Count of IDA-1 puncta in the synaptic region defined as 50-70 μm region that follows 45 μm asynaptic region from the turn of the commissure. (H-M) Alignments of 5 axons expressing INS-22∷Emerald in *ebp-1; ebp-2* double mutants per rescue transgene. (N) Quantification of INS-22∷Emerald fluorescence in *ebp-1* mutant with or without respective rescue transgene. (O) Quantification of INS-2∷Emerald fluorescence in *ebp-1; ebp-2* mutant with or without respective rescue transgene. n=18-38 ****p<0.0001; *p<0.05 (one-way ANOVA). CH, Calponin homology domain. CC, coiled-coil domain. EBH, End-binding Homology domain. FL, full length.

To complement these loss of function experiments, we also tested the sufficiency of EBP-1 domains and EBP-1 homologs for rescuing *ebp-1* mutants. DA9 specific expression of EBP-1 or its mammalian homolog EB1 fully rescued the loss of INS-22 labelled DCVs in *ebp-1* mutants, indicating that EBP-1 functions cell autonomously and that EB1 function is conserved from invertebrates to vertebrates (**Figure 2N**). EBP-2 overexpression also rescued *ebp-1* mutants (**Figure 2N**), suggesting that at high expression levels the functional differences between these proteins can be overcome. We next expressed either the Calponin Homology (CH) domain, which mediates MT tip tracking, or the remaining EBP-1 C-terminus comprising the EBH and acidic tail. These experiments were done *ebp-1; ebp-2* double mutants to prevent potential dimerization between EBP-1 fragments and native EBP-2.^33^ Interestingly, the CH domain could not rescue the double mutant phenotype, whereas the remaining C-terminus was sufficient to restore INS-22 levels to the axon (**Figure 2J** and **2K**). The Y264A and E272A mutations which interfere with EB binding to SxIP motifs did not prevent the C-terminus from rescuing *ebp-1; ebp-2* double mutants (**Figure 2K**). In contrast, a construct lacking both the EBH domain and acidic tail could not rescue the phenotype (**Figure 2L**). Finally, we found that the acidic tail was not required for rescuing activity and that expressing the EBH domain on its own could restore the INS-22 fluorescent signal to the axon of double mutants (**Figure 2M** and **2O**). These results highlight the importance of the EBH domain in ensuring axonal DCV delivery.

### EBP-1 localizes near AP-1μ positive Golgi carriers and is required for cell body exit of DCVs

Since loss of DCVs from the axon in *ebp-1* mutants was not due to increased secretion, we next tested whether it reflected reduced cargo delivery from the cell body to the axon. For this, we measured how many endoIDA-1/Phogrin positive vesicles exited the cell body after photobleaching the residual signal of endoIDA-1 in the proximal axon (**Figure 3A**). In *ebp-1* mutants, we observed a much higher incidence of zero exit events (no DCV exiting the cell body during an imaging window) over multiple experimental replicates (**Figure 3D** and **3E**), suggesting that EBP-1 promotes cell body exit of DCVs. The velocity of endoIDA-1 puncta exiting the cell body was similar between wildtype and mutants (**Figure 3B** and **3C**), indicating that EBP-1 is not required for UNC-104/KIF1A motility in the proximal axon. These results raise the hypothesis that reduced cell-body exit underlies the reduced levels of axonal DCVs in *ebp-1* mutants.

**Figure 3:**
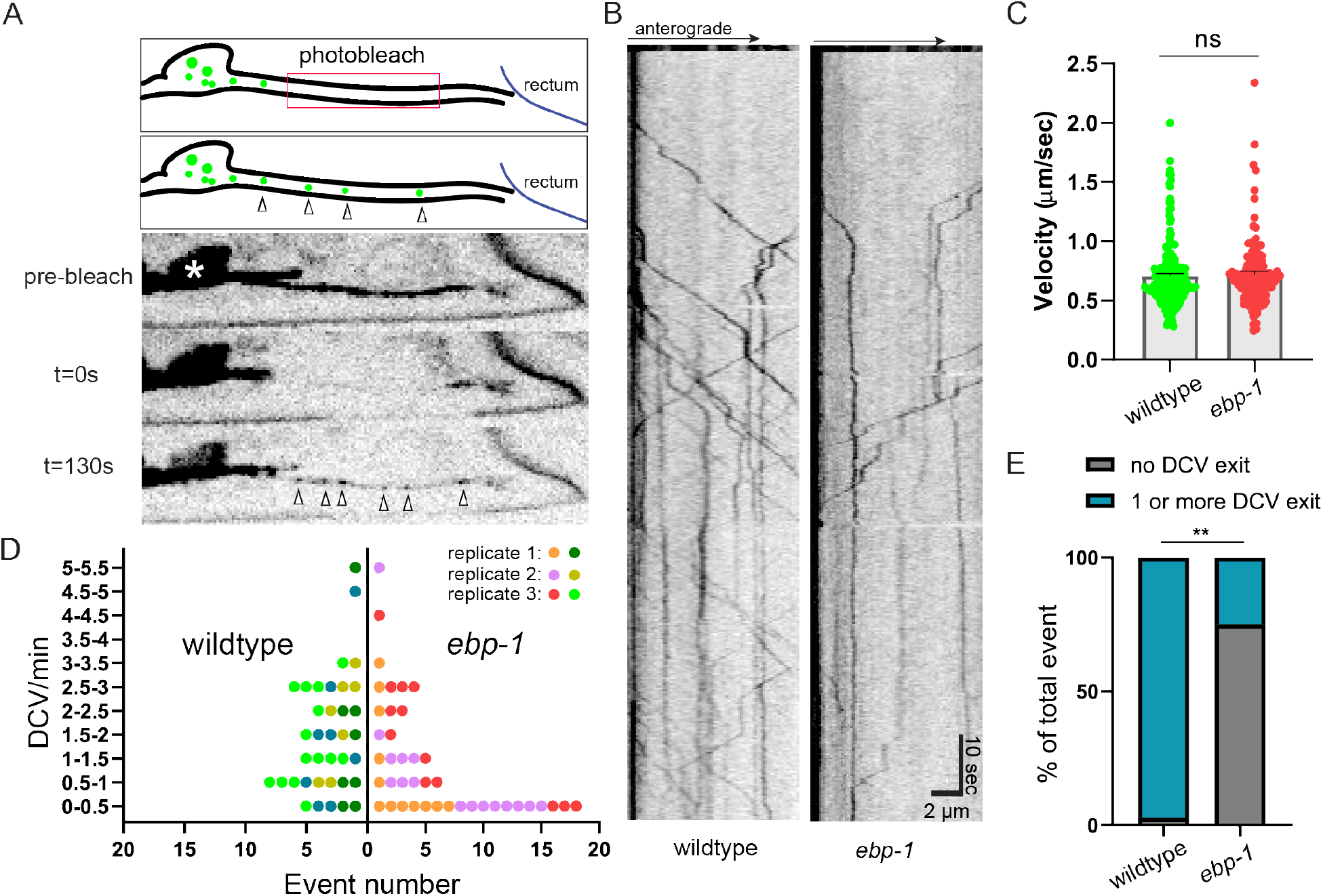
*ebp-1* is required for efficient cell body exit of DCVs. (A) Three frame montage of photobleaching experiment with diagram. Proximal axon outside the soma (white asterisk) is bleached to eliminate background from residual DCVs and allow visualization of DCVs exiting the cell body (indicated with empty arrow heads). Cell body exit events of endoIDA-1 positive DCV are imaged for 120-180 seconds at three frames/second. (B) Kymograph of endogenous DCV exiting cell body in wildtype and *ebp-1* mutant. Scale bar is ten seconds vertical and 2 μm horizontal. (C) Velocity of DCV exiting the cell body in wildtype and *ebp-1* mutant. n=133-183 events in 27-31 animals. t-test (D) “Superplot” of exit event frequency over multiple replicates.^63^ Each dot indicates an animal and each color indicates different experimental replicates. Wildtype is shown in green color palette and *ebp-1* mutant in red. n=37-41. (E) Proportion of zero event (no DCV exiting during imaging window) in wildtype and *ebp-1* mutant. **p<0.01 (n is same as (D), Chi-square test).

To examine the potential function of EBP-1 in the cell body, we examined the subcellular localization of endogenously tagged EBP-1 and EBP-2. We tagged both proteins at the C terminus and confirmed that they were functional by MT tip tracking (**Figure S3D**), consistent with previous studies using tagged EBs.^49^ Both EBP-1 and EBP-2 were uniformly distributed throughout the neurons as expected for a cytosolic protein (**Figure S3A** and **S3B**). Interestingly, in the cell body, a local accumulation of EBP-1 was observed in two or three punctate structures (**Figure 4B**). This localization was not unique to DA9 but could be observed across many neuronal cell bodies in the ventral cord (**Figure 4A**). EBP-2 did not show such clear localization to cell body puncta, either in a wildtype background or in *ebp-1* mutants (**Figure 4B and S3C**).

**Figure 4:**
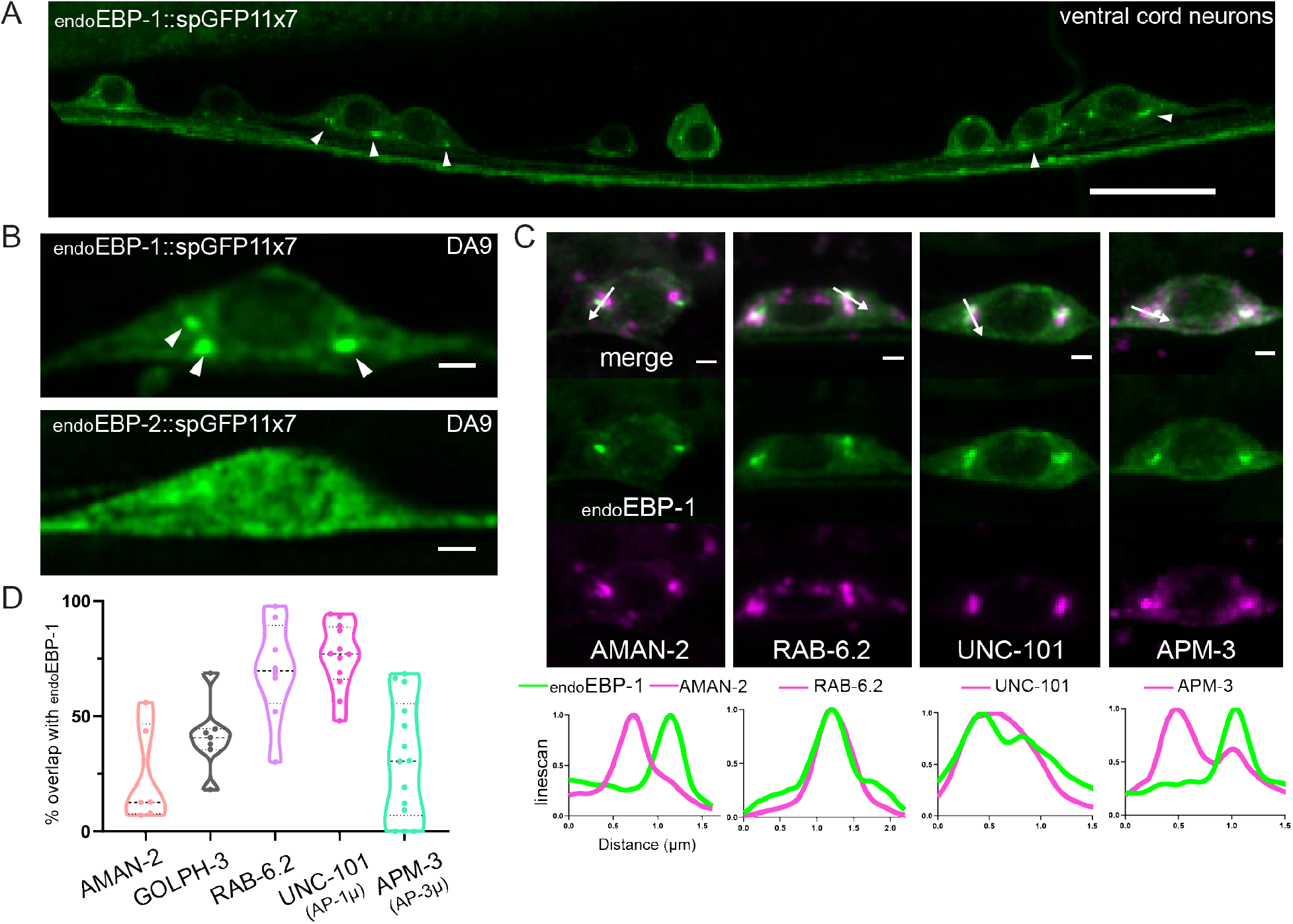
Endogenous EBP-1 is enriched in the vicinity of late Golgi carriers. (A) Airyscan image of endogenous EBP-1 in ventral neurons that express *Pmig-13∷spGFP1-10*. Note consistent enrichment of EBP-1 in 2-3 puncta per neuronal cell body indicated by white arrowheads. Scale bar 10 μm. (B) Airyscan images of endogenous EBP-1 and EBP-2 in the DA9 cell body. White arrowheads pointing EBP-1 cell body structure. Scale bar 1 μm. (C) Endogenous EBP-1 in green and various secretory pathway markers in magenta, imaged in DA9 cell body. A linescan is drawn in the direction of the white arrow. Scale bar 1 μm. (D) Quantification of precent area overlap between endogenous EBP-1 and each Golgi marker in maximum projection image of DA9 cell body z-stack.

EBP-1 puncta in the cell body were highly reminiscent of Golgi stacks in *C. elegans* neurons, prompting us to test colocalization with Golgi markers.^15, 50^ Using Airyscan microscopy, we observed that AMAN-2, a cis-medial Golgi marker, did not show high overlap with EBP-1, whereas the trans Golgi marker GOLPH-3 had a higher overlap (**Figure 4C** and **4D**). Interestingly, EBP-1 showed 69.7% overlap with a trans Golgi/late Golgi vesicular carrier marker, RAB-6.2. This suggests that EBP-1 localizes in the vicinity of the trans Golgi and post Golgi carriers. Previous work has shown that different subdomains of the trans Golgi show a differential enrichment of the cargo adaptor complexes AP-1 and AP-3.^14^ AP-3μ /APM-3 is important for sorting SVPs to the axon while AP-1μ/UNC-101 is required for sorting DCV cargo.^14, 20^ We found that EBP-1 had an average of 79.1% overlap in area with UNC-101, while it only overlapped 31.2% with APM-3 (**Figure 4C** and **4D**). Together, these results indicate that EBP-1 is required for cell body exit of endoIDA-1 and is enriched in cell body puncta in the vicinity of trans Golgi carriers and the DCV sorting machinery AP-1μ/UNC-101.

### EBP-1 functions in the neuronal soma to promote DCV trafficking to the axon

The enrichment of EBP-1 in cell body puncta and its role in promoting cell body exit of DCVs suggested that the neuronal soma might be the site where it functions in DCV trafficking. We tested whether EBP-1 co-traffics with endoIDA-1 or INS-22 but could not detect co-movement events in the axon (not shown), suggesting that any interaction with exiting DCVs may be transient or that EBP-1 may function prior to DCVs entering the axon. To test whether EBP-1 functions in the cell body prior to DCV exit, we sought to restrict its location by fusing it to the integral trans Golgi proteins BRE-4 or GOLGIN-84. For this chimera, we used a version of EBP-1 lacking the CH domain to prevent interactions with MTs from affecting the localization of the chimeric protein and since expression of EBP-1 lacking this domain could rescue *ebp-1; ebp-2* mutants (**Figure 2K**). Both RFP∷EBP-1ΔCH∷BRE-4 and RFP∷EBP-1ΔCH∷GOLGIN-84 chimeras showed punctate localization in the cell body and could not be detected in the axon (**Figure 5D’ and 5D’’**). Interestingly, this chimeric protein was sufficient to restore endoIDA-1 to the axon in *ebp-1* mutants (**Figure 5E**). Control constructs RFP∷BRE-4 or RFP∷EBP-1ΔCHΔC∷BRE-4 did not rescue axonal DCVs, consistent with the inability of overexpressed EBP-1ΔCHΔC to rescue *ebp-1; ebp-2* mutants (**Figure 5F, 5H**, and **5I**). These data support the conclusion that EBP-1 promotes DCV trafficking to the axon by acting in the vicinity of the Golgi in the cell body.

**Figure 5:**
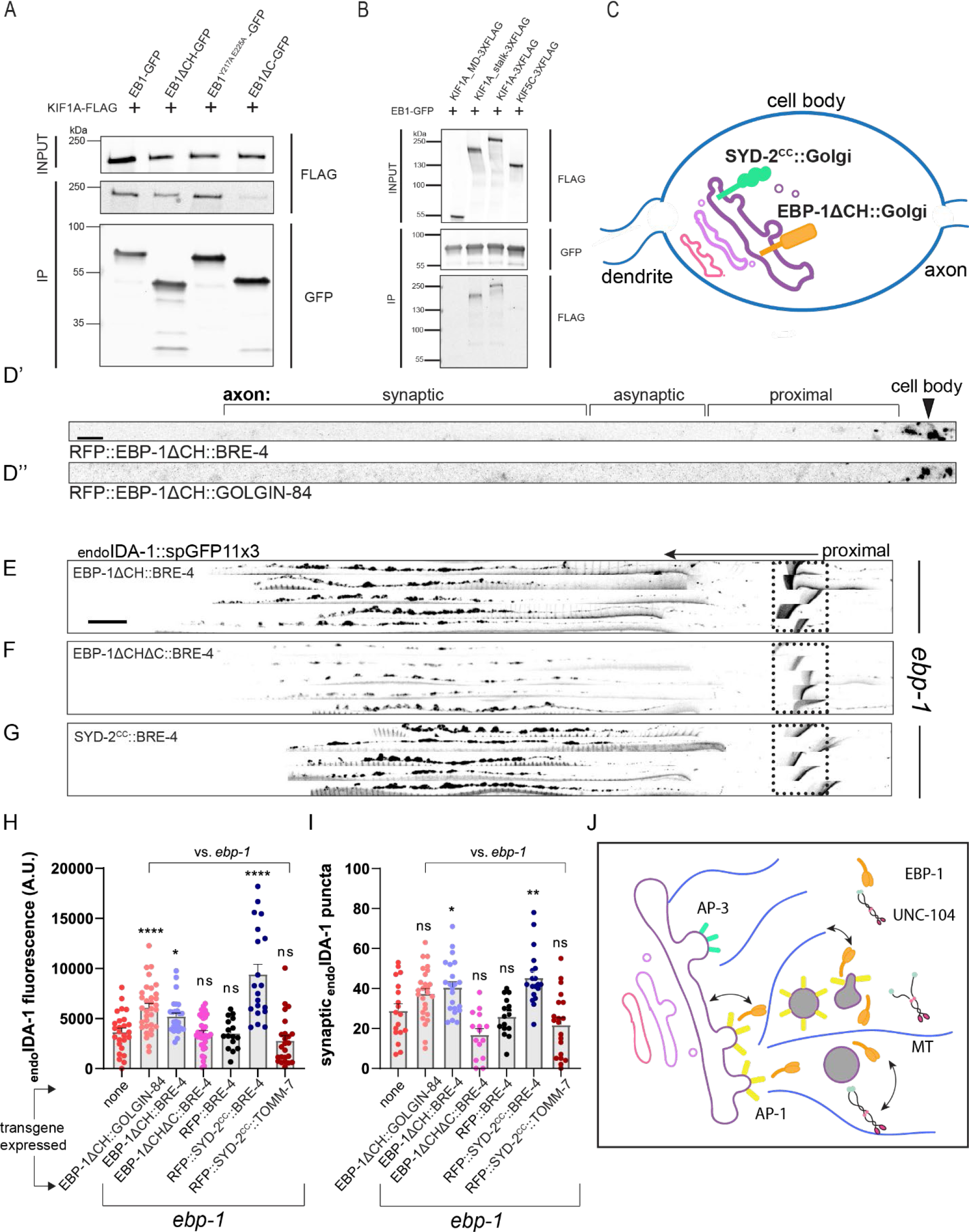
EB1 interacts with KIF1A and EBP-1 function in neuronal soma to promote DCV trafficking. (A) co-IP between EB1 fragments and KIF1A. EB1 fragments stained with anti-GFP and KIF1A with anti-FLAG. (B) co-IP between EB1 and KIF1A motor domain, KIF1A stalk, KIF1A, and KIF5C. (C) Diagram of Golgi protein chimera with EBP-1ΔCH and SYD-2^CC^. (D) Representative images of RFP∷EBP-1ΔCH∷BRE-4 (D’) and RFP∷EBP-1ΔCH∷GOLGIN-84 (D’’) showing that the chimeric proteins are restricted to the DA9 cell body. Black arrowhead indicates DA9 cell body. Scale bar 10 μm. (E-G) Alignments of 5 representative axons per rescue showing endoIDA-1∷GFP in DA9. Black arrow points distally. Autofluorescence from rectum in dotted box. Scale bar 10 μm. (H) Quantification of fluorescence of endoIDA-1 for each rescue condition. n=16-34. (I) Count of endoIDA-1 puncta in the synaptic region. n=15-26. (J) Model diagram showing EBP-1 biasing kinesin-3 localization by interaction or binding microtubule near Golgi. ****p<0.0001; **p<0.01, *p<0.05 (one-way ANOVA for (F) and (G). t-test for (H) and (I)).

### EB1 interacts with KIF1A and a KIF1A/UNC-104 interacting domain can replace EBP-1 in the soma

Since the DCV phenotype of *ebp-1* mutants was suggestive of transport defects, we tested whether it could involve UNC-104/kinesin-3, which is the motor for DCVs. Previously, EBs have been shown to physically interact with other kinesin motors and with dynactin, either directly or indirectly.^27, 51–55^ However, an interaction between KIF1A/UNC-104 and EB has not been described. *C.elegans* UNC-104 formed aggregates when expressed in mammalian cell culture (not shown), so we tested its highly conserved mammalian homolog KIF1A and the mammalian EB1, which could robustly rescue the DCV phenotype of *ebp-1* mutants in *C. elegans* (**Figure 2N**). FLAG tagged KIF1A and GFP tagged EB1 co-immunoprecipitated from HEK293T cells, while the neuronally enriched kinesin-1/KIF5C did not co-immunoprecipitate with EB1 (**Figure 5B fourth lane**). EB1 without the Calponin Homology domain (EB1ΔCH) was also able to pull down full length KIF1A (**Figure 5A** second lane), albeit not as efficiently as full length EB1, suggesting that MT binding could facilitate this interaction (**Figure 5A**). Interestingly, the EB1ΔC construct minimally co-immunoprecipitated with KIF1A but EB1^Y217A E225A^ -GFP still pulled down KIF1A (**Figure 5A** third and fourth lane). We also found that the motor domain of KIF1A was dispensable for this interaction and the stalk region alone could co-immunoprecipitate with EB1-GFP (**Figure 5B** first and second lane), consistent with previous reports of EBs interacting with other kinesins via their stalk regions.^51, 53–55^

We noted that the domains that mediated the EB1-KIF1A interaction reflected the ability of DA9-expressed EBP-1 fragments to rescue the DCV phenotype of *ebp-1; ebp-2* mutants (**Figure 2K** and **2O**), suggesting that an interaction between EBP-1 and UNC-104/KIF1A might be important for DCV trafficking. Since we found that EBP-1 functions in the cell body near Golgi-derived carriers, we asked whether it acts by transiently recruiting UNC-104 to these sites. For this, we tested whether *ebp-1* mutants can be rescued by artificially recruiting UNC-104/KIF1A to the Golgi with a chimera between the Golgi-resident protein BRE-4 and a UNC-104/KIF1A binding domain from a protein not related to EBs. We used the UNC-104 interacting coiled-coil domain from SYD-2/liprin-α (SYD-2^coiled-coil^), which was shown to directly interact with kinesin-3 in worms and mammals.^56, 57^ RFP∷SYD-2^coiled-coil^∷BRE-4 expression in DA9 was able to fully rescue the loss of endoIDA-1 from the axon of *ebp-1* mutants (**Figure 5G, 5H**, and **5I**). In contrast, RFP∷SYD-2^coiled-coil^∷TOMM-7, which localized to mitochondria in the cell body and axon, could not rescue (**Figure 5H** and **5I**). This result suggests that transiently localizing UNC-104 to the vicinity of Golgi carriers is sufficient to compensate for EBP-1 function in promoting axonal delivery of DCVs. Collectively, our results support a model in which Golgi-localized EBP-1 promotes axonal DCV transport by facilitating motor-cargo association through its interactions with kinesin-3/UNC-104 and with microtubules (Figure 5J).

## Discussion

How specific motor-cargo associations are regulated in the cell body prior to long distance transport in the axon is unclear. Here we report a role for End Binding protein in promoting the initial steps of transport of one KIF1A/UNC-104 cargo – dense core vesicles (DCVs). Loss of EBP-1 leads to a specific reduction in axonal DCVs, while other cargoes (ATG-9, RAB-3, SNG-1) remain unperturbed. This reduction can be traced back to a reduced number of DCVs exiting the cell body, suggesting that neuronal soma is where EBP-1 functions. Consistently, EBP-1 (but not its close paralog EBP-2), is enriched in the cell body at the Golgi complex in the vicinity of AP-1μ which sorts DCVs at the trans Golgi. EB1 co-immunoprecipitates with KIF1A in CH and EBH domain dependent manner and overexpression of its EBH domain is sufficient to rescue *ebp-1* mutants. Importantly, loss of DCVs from the axon can be rescued by tethering a MT-binding deficient version of EBP-1 to the Golgi or by anchoring to this organelle a KIF1A/UNC-104 interacting domain from an unrelated protein. Together, these results support a model in which EBP-1 promotes the recruitment of UNC-104 to nascent DCVs at the trans Golgi by using its ability to bind both the motor and the microtubules in order to facilitate the initial steps of axonal transport. These results provide insight into how the specificity of motor-cargo association in the cell body is generated and reveal an unexpected role of a MT associated protein.

### Coupling cargo sorting with motor-association in the neuronal soma for efficient axonal transport

In neurons, the biogenesis of SVPs and DCVs involves complex sorting steps, followed by an association between the transport packet and the axonal motor, but how these processes are coordinated is unclear. ^15, 58, 59^ Particularly in cases where the same motor carries different cargo, specific mechanisms that promote its association with given cargos should exist. KIF1A/UNC-104 has a Pleckstrin Homology domain at its C-terminal tail that has been shown to bind PI(4,5)P2 and PI(4)P and is thought to be the main determinant in cargo recognition.^3–5, 17^ The importance of the PH domain was shown in mammalian neurons and in *C. elegans* where deletion or a single point mutation in the PH domain abolishes trafficking of DCVs, SVPs, lysosomes, and ATG-9 vesicles.^3, 5, 17^ However, it is unclear if membrane recognition by a PH domain is sufficient to explain the transport of vesicles that likely differ in size and membrane properties or to support differential delivery of these different cargo into the axon.

We found that EBP-1 specifically promotes the cell body exit of DCVs transported by UNC-104. EBP-1, but not its close homolog EBP-2, is specifically enriched near the adaptor complex that sorts DCVs, but not SVPs, for axonal delivery in DA9. Thus, we propose that spatial segregation between these adaptor complexes could underlie the ability of EBP-1 to specifically promote association between UNC-104 and DCVs. Additional factors likely exist that promote UNC-104 association with other cargo.^17, 60^ How EBP-1 is specifically enriched near AP-1μ is unclear. In non-neuronal cell culture, a complex of GM130 / AKAP450 / Myomegalin promotes EB1/3 localization to Golgi.^29^ These proteins (including EB1/3) are required for normal Golgi architecture and MT organization, making it challenging to precisely dissect roles in promoting transport. We did not observe gross Golgi abnormalities at the level of light microscopy in DA9 in *ebp-1* mutants, suggesting that they do not play a structural role there.

### Neuronal function of EBs

EBs are highly conserved, and while they are routinely used as markers of growing MT ends, the full range of their physiological roles remains to be discovered. This observation is underscored by the surprisingly mild phenotype in mammalian cells where all 3 EBs are disrupted or in worm embryos with a triple EBP deletion.^29, 33, 34^ In neurons, EBs were proposed to organize neuronal MTs by guiding polymer growth along other MTs, linking MTs to actin-based structures or to the AIS, or controlling polymer dynamicity.^26–28, 31, 32^. We show that in addition to these cytoskeletal functions, which mostly take place along neurites, EBP-1 has a specific role in promoting the transport of DCVs from the cell body by UNC-104/KIF1A. This EB function is conserved since EB1 could rescue *ebp-1* mutants and interacted with KIF1A.

The localization of EBs to MT ends makes them ideal for regulating molecular motors, which often associate with the plus ends, or fall from it, at the end of a run.^30^ Consistently, EBs were shown to interact with several kinesins and to help load dynactin/dynein on MTs at the distal axon.^32, 51, 53–55^. In addition, EB on the Golgi in non-neuronal cells can recruit microtubules to the Golgi.^29^ Together with our data our results showing that EBP-1 acts at the Golgi through UNC-104/KIF1A, we propose that EBP-1 enriches the motor near DCV sorting sites by interacting both with the motor and with the microtubule. This model is consistent with the necessity we observe for the CH domain in CRISPR mutants and with the ability of EBH domain to rescue *ebp-1* mutants when overexpressed.

A second site where EBP-1 one might act through UNC-104/KIF1A is presynapses. We observed reduced spacing of presynaptic sites in *ebp-1* mutants, which is the opposite phenotype of *unc-104* gain of function alleles.^1^ Furthermore, EB1 competes with the motor domain of KIF1A for GTP-cap binding in vitro, and this was proposed to facilitate cargo unloading at presynaptic sites of cultured hippocampal neurons, where MTs are highly dynamic.^30, 61^ In DA9 it is possible to separate the cell body and synaptic functions of EBP-1: the former is not affected by deleting only the CH domain and can be rescued from the cell body, whereas the latter strongly depends on the CH domain and cannot be rescued from the cell body (Figure S4A).

The ability to assess EB function using CRISPR editing of endogenous loci provides an opportunity to dissect functional differences between EB family members. For example, we found that both EBP-1 and EBP-2 are expressed at similar levels in DA9, but EBP-1 plays a more significant role in DCV transport. Despite their overall similarity, only EBP-1 is enriched at the Golgi, while EBP-2 is significantly more efficient than EBP-1 in tracking growing MT plus-ends. These observations mirror results from other systems where differential microtubule binding preferences has been observed between EB paralogs owing to structural differences of each calponin homology domain.^62^. Importantly, the differences between EBs can sometimes be overcome by overexpression: EBP-2 overexpression rescue *ebp-1* mutants. Similarly, EBH domain deletion does not phenocopy *ebp-1* deletion, but its ectopic expression is sufficient to restore DCV transport in *ebp-1; ebp-2* double mutants. These results likely reflect the existence of redundant protein networks at the MT tip, which are supported both by interacting proteins binding each other as well as associating with the MT polymer. Future studies will be required to precisely determine the sequences that confer specific subcellular localizations to EBs or that dictate their tip-tracking efficiency, which should also lead to a better understanding of their physiological roles.

## Acknowledgments

The authors thank the CGC and NBRP Mitani lab for various strains. We also thank Kang Shen for strains and Sander van den Heuvel for SV1877 strain. We would also like to thank Michael Koelle for LX2519 strain as well as sharing resources for egg-laying behavior and Selim Çetinkaya for the initial worm husbandry for the project. We thank the members of the Yogev, De Camilli, and Hammarlund labs for discussions and technical assistance. This work was supported by NIH R35-GM133573 and DK45735.

## Author contributions

J.P, P.D.C, and S.Y designed the project, carried out experiments in *C. elegans* and wrote the manuscript. K.G shared the initial discovery with *ebp-1* mutant animals and DCV marker *ceIs308*.

---

**Supplemental Figure 1:**
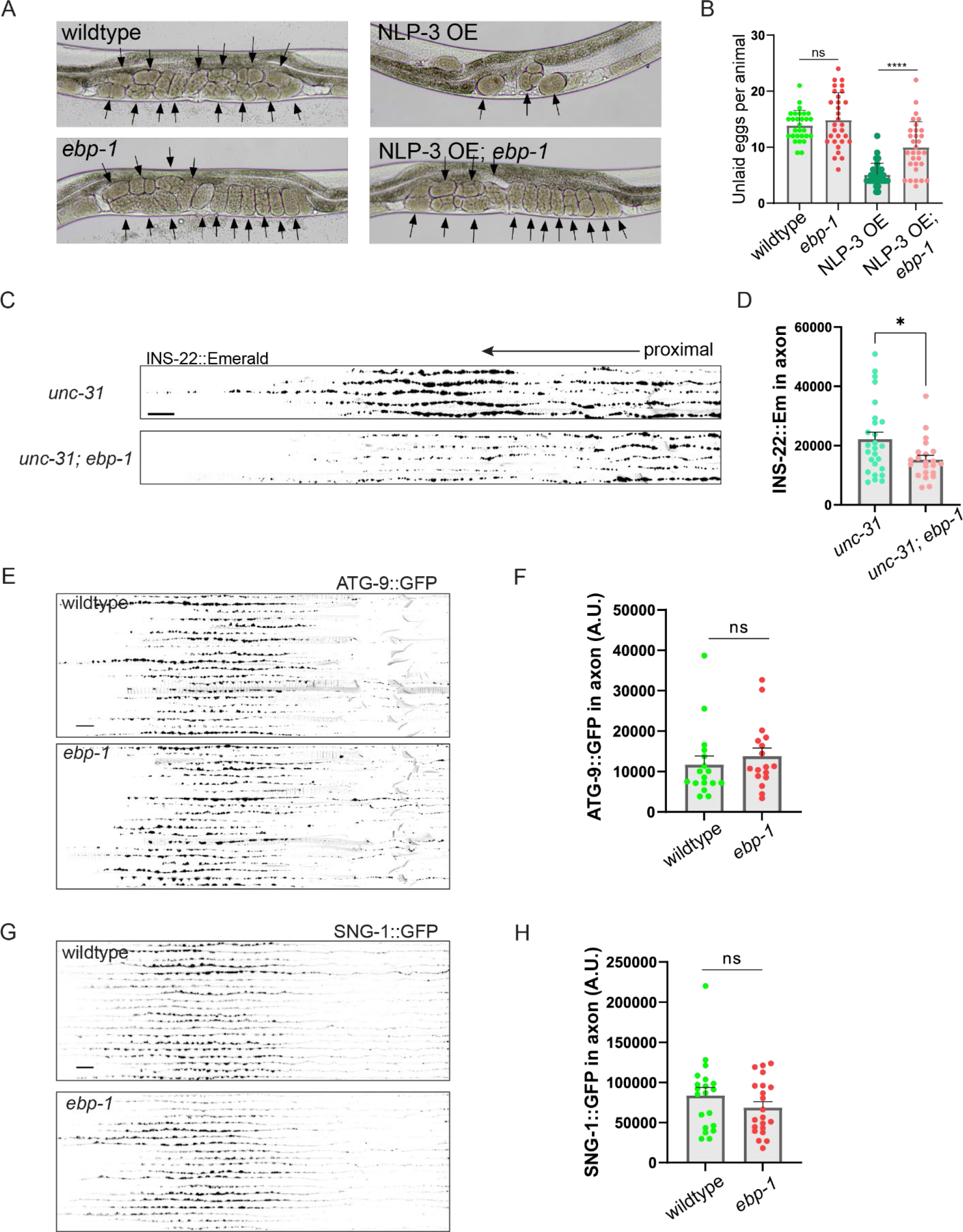
*ebp-1* is required for the axonal delivery of DCVs, not SVPs. (A) Images of worm uterus with eggs inside. Eggs are indicated by the black arrowheads. (B) Count of unlaid eggs in uterus of synchronized adults. n=29-32 (C) Alignments of 5 representative axons per genotype showing INS-22∷Emerald in DA9. Black arrow points distally. (D) Quantification of fluorescence of INS-22∷Emerald. (E, G) Alignments of 20 representative axons per genotype showing ATG-9∷GFP (E), and SNG-1∷GFP in DA9 (G). (F) Quantification of ATG-9∷GFP fluorescence. (H) Quantification of SNG-1∷GFP fluorescence. n=17-26. Scale bar 10 μm for all scale bars shown. ****p<0.0001; *p<0.05 (one-way ANOVA for (B). t-test for (D), (F), (H))

**Supplemental Figure 2:**
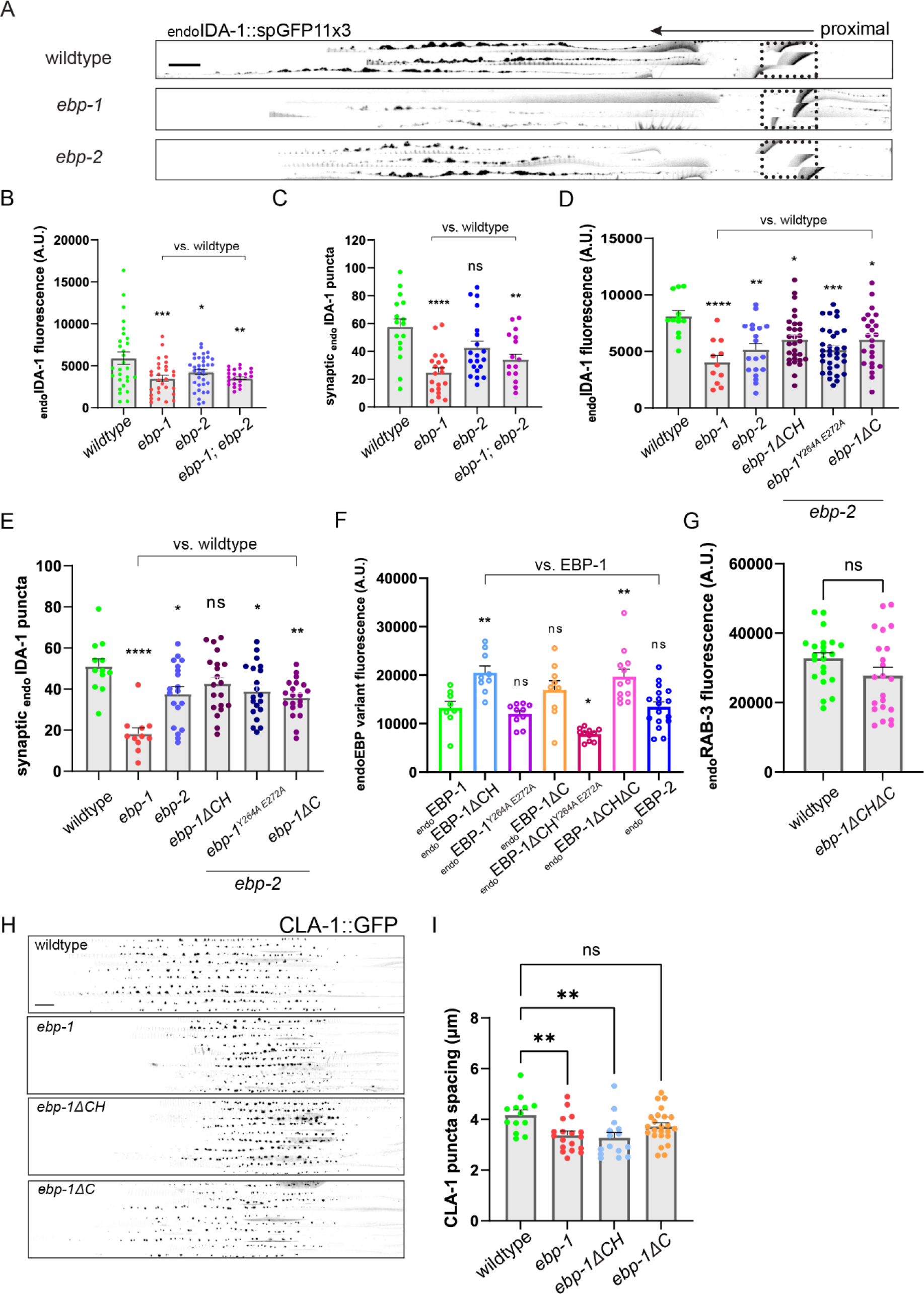
EBP-1 truncations are stably expressed and ebp-1 mutants show reduced synaptic spacing. (A) Alignments of 3 representative axons per genotype showing endoIDA-1∷GFP in DA9. Black arrow points distally. Autofluorescence from rectum in dotted box. Scale bar 10 μm. (B, D) Quantification of fluorescence of endoIDA-1∷GFP. n=11-35 (C, E) Count of endoIDA-1 puncta in the synaptic region. n=11-20. (F) Quantification of endoEBP-1 truncations and endoEBP-2 fluorescence in DA9. n=8-16 (G) Quantification of endogenous RAB-3 fluorescence in DA9. n=22 per genotype. (H) Alignments of 20 representative axons per genotype showing CLA-1∷GFP in DA9 synaptic region. Scale bar 10 μm. (I) Spacing measure between each CLA-1∷GFP puncta in μm. n=8-35. ****p<0.0001; ***p<0.001; **p<0.01, *p<0.05 (one-way ANOVA for (B), (C), (D), (E), (F), and (I). t-test for (G)).

**Supplemental Figure 3:**
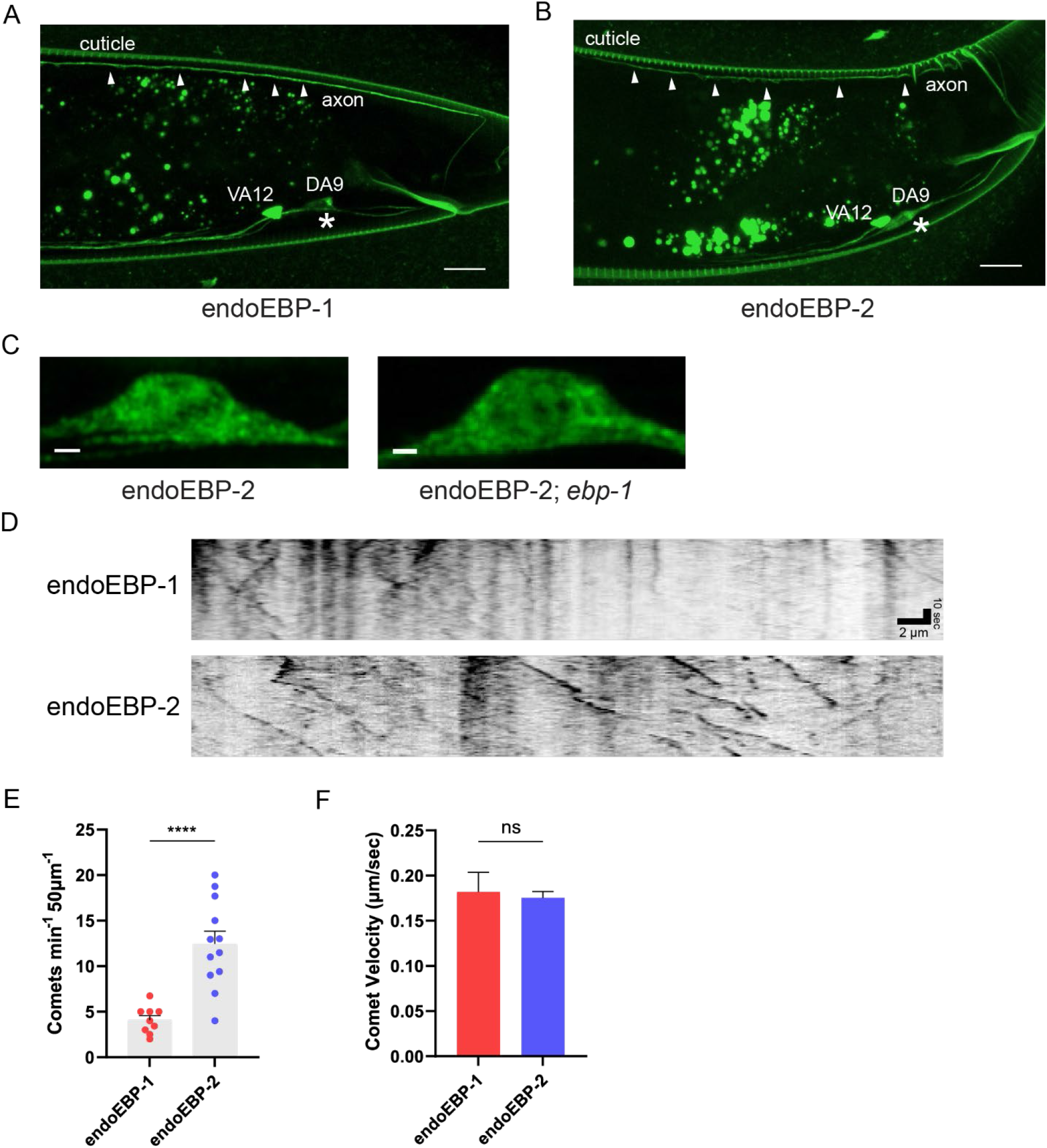
endoEBP-1 and endoEBP-2 localization and microtubule tracking behavior. (A, B) Representative image of endoEBP-1 and endoEBP-2 distribution in DA9. endoEBP-1 and endoEBP-2 signal shown right above white arrowheads. Scale bar 10 μm. (C) Airyscan image of endoEBP-2 in wildtype and *ebp-1* mutant. Scale bar 1 μm. (D) Representative kymograph of endoEBP-1 and endoEBP-2 comets in ALNL neuron. Scale bar is ten seconds vertical and 2 μm horizontal. (E) Quantification of comet frequencies and (F) velocity of endoEBP-1 and endoEBP-2. n=45-154 comet events from 9-12 animals.

**Supplemental Figure 4:**
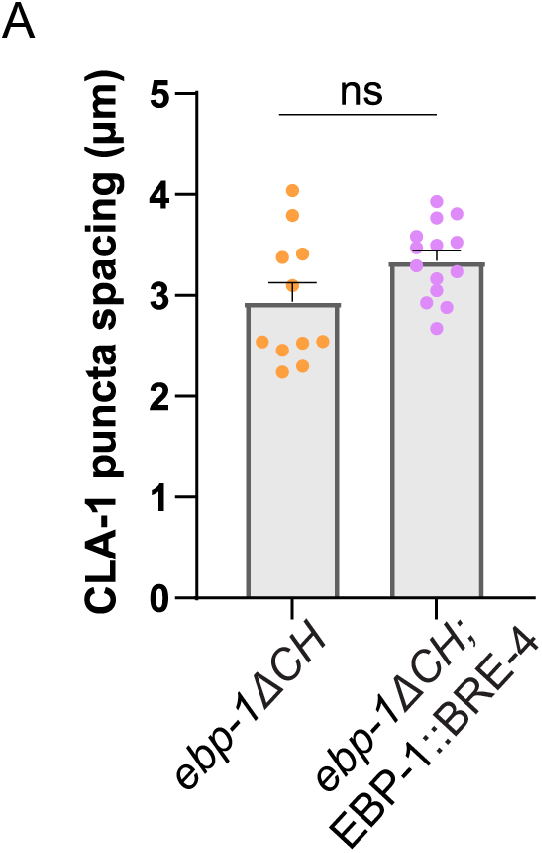
Presynapse spacing defect cannot be rescued by cell-body restricted EBP-1. (A) Spacing measure between each CLA-1∷GFP puncta in μm. Comparison between *ebp-1ΔCH* mutant with or without EBP-1∷BRE-4 rescue array. n=11-34. t-test.

## RESOURCE AVAILABILITY

### Lead contact

Further information and requests for reagents should be directed to the lead contact.

### Materials Availability

All transgenic *C. elegans* strains generated for this study are available from the lead contact upon request.

### Data and Code Availability

Code and raw data used for this study are available from the lead contact upon request.

## EXPERIMENTAL MODEL AND SUBJECT DETAILS

### Strains and maintenance

All *C. elegans* strains were grown on nematode growth medium plates seeded with *E. coli* OP50 as previously described.^64^The N2 Bristol strain was used as wildtype and for outcrosses. All animals were grown at 20°C for experiments/injection.

## METHOD DETAILS

### Cloning and constructs

Genes of interest were PCR cloned from either cDNA or gDNA of N2 Bristol strain using PfuUltra II Fusion HS DNA polymerase (Agilent). Amplicon was ligated into pSM vector backbone for expression in *C. elegans* and EGFP-C1 (Clontech) for mammalian cells. All plasmid ligation step was done using In-fusion seamless cloning (Takara Bio). Appropriate mutagenesis was done using QuikChange Lightning Multi Site-Directed mutagenesis kit (Agilent). Final product plasmids were confirmed by Sanger sequencing. All plasmids are freely available upon request.

### CRISPR/Cas9 genome editing

All CRISPR strains were generated following the published protocol.^65^ Custom designed crRNA, trRNA, and Cas9 are from IDT. Details of crRNA sites and repair template used is in the supplemental table 2.

### Fluorescence microscopy and sample preparation

L4 larvae animals were grown at 20°C, 20-24 hours prior to imaging. One day adult animals were used mainly for quantification and assessment of phenotype. For endogenous EBP-1/EBP-2 comet imaging, adult animals were bleached and synchronized L1 larvae were isolated to image L3 worms. For general imaging, worms were immobilized on agar pads with 10 mM Levamisole dissolved in M9 buffer. For time-lapse imaging, one day adult animals were incubated in 0.5 mM levamisole at room temperature for 6 min 30 seconds (4 min 30 seconds for L3) prior to mounting on 10% agarose pad in M9 without additional paralytics. Andor Dragonfly spinning-disk confocal microscope equipped with a plan apochromat objective (63x, 1.4 NA, oil) and a Zyla scientific CMOS camera was used for standard fluorescence imaging. The same imaging settings (laser power, exposure time, and gain) were used throughout the study.

In photobleaching experiments, region of interest was photobleached using 405 nm laser with a Perkin-Elmer Photokinesis FRAP unit. Pre-bleach and post-bleach time lapse was imaged at 3 Hz using an Olympus BX61 microscope equipped with a Hamamatsu ORCA-Flash4.0 LT camera. For high resolution imaging, Airyscan function of Carl Zeiss LSM880 confocal laser scanning microscope (63x, 1.4 NA, plan-apochromat, oil) with Airyscan detector was used. Raw Airyscan images were processed using ZEN imaging software (Zeiss).

### Egg laying quantification

Quantification of unlaid egg was performed using adult animals 30 hours after staging late L4 larvae. The adult animals were bleached to dissolve the body and impermeable eggs were counted on a 96-well plate. Precise protocol is detailed in Brewer et al., 2019.

### Cell culture, transfection, and co-IP

HEK293T cells (American Type Culture Collection) were cultured in Dulbecco’s Modified Eagle Medium (DMEM) containing 10% Fetal Bovine Serum, 2 mM L-glutamine, 1 mM sodium pyruvate, 100 U/mL penicillin, and 100 mg/mL streptomycin (Gibco) at 37 °C and 5% CO2. For transfection, equimolar amount of EB1-GFP, EB1 fragment-GFP, and KIF1A-3XFLAG were transiently transfected using FuGene HD (Promega)/Lipofectamine 2000 (Thermo Fisher). 48 hours after transfection, cells were lysed with a buffer containing 1% Triton X-100 for 20 min and the lysate was incubated with anti-GFP magnetic beads (Chromotek) for 2 hours in 0.33% Triton X-100 solution. After incubation, beads were washed and the samples were eluted with Sodium dodecyl sulfate-polyacrylamide gel electrophoresis (SDS-PAGE) sample buffer.

Samples were then run on 4~20% Mini-PROTEAN TGX Stain-Free gel and transferred to 0.2 μm nitrocellulose membrane (both Bio Rad). Membrane was incubated with M2 anti-FLAG (Sigma-Aldrich) and anti-GFP (Abcam) overnight and after few washes was stained with Li-COR fluorescence secondary antibodies (Li-COR) for imaging.

## QUANTIFICATION AND STATISTICAL ANALYSIS

### Fluorescence quantification

Raw imaging files were imported to ImageJ for further analysis. Puncta number was counted using FindFoci plugin using similar settings for all images analyzed.^66^ Kymographs were generated with ImageJ Multi Kymograph plugin where resulting traces were analyzed manually in Microsoft Excel. Movement correction was done with StackReg plugin for time-lapse movies with minor drift before kymograph analysis.

### Statistical analysis

Statistical analysis was performed on Prism 9 (GraphPad) and Microsoft Excel. Three experimental replicates were included in each data set and data were considered significant at *p* ≤ 0.050 with statistical test indicated in each figure legend.

## References

1. Niwa, S., Lipton, D.M., Morikawa, M., Zhao, C., Hirokawa, N., Lu, H., and Shen, K. (2016). Autoinhibition of a Neuronal Kinesin UNC-104/KIF1A Regulates the Size and Density of Synapses. Cell Rep 16, 2129–2141.

2. Hammond, J.W., Cai, D., Blasius, T.L., Li, Z., Jiang, Y., Jih, G.T., Meyhofer, E., and Verhey, K.J. (2009). Mammalian Kinesin-3 motors are dimeric in vivo and move by processive motility upon release of autoinhibition. PLoS Biol 7, e72.

3. Klopfenstein, D.R., Tomishige, M., Stuurman, N., and Vale, R.D. (2002). Role of Phosphatidylinositol(4,5)bisphosphate Organization in Membrane Transport by the Unc104 Kinesin Motor. Cell 109, 347–358.

4. Kumar, J., Choudhary, B.C., Metpally, R., Zheng, Q., Nonet, M.L., Ramanathan, S., Klopfenstein, D.R., and Koushika, S.P. (2010). The Caenorhabditis elegans Kinesin-3 motor UNC-104/KIF1A is degraded upon loss of specific binding to cargo. PLoS Genet 6, e1001200.

5. Klopfenstein, D.R., and Vale, R.D. (2004). The lipid binding pleckstrin homology domain in UNC-104 kinesin is necessary for synaptic vesicle transport in Caenorhabditis elegans. Mol Biol Cell 15, 3729–3739.

6. Hall, D.H., and Hedgecock, E.M. (1991). Kinesin-related gene unc-104 is required for axonal transport of synaptic vesicles in C. elegans. Cell 65, 837–847.

7. Stavoe, A.K., Hill, S.E., Hall, D.H., and Colon-Ramos, D.A. (2016). KIF1A/UNC-104 Transports ATG-9 to Regulate Neurodevelopment and Autophagy at Synapses. Dev Cell 38, 171–185.

8. Zahn, T.R., Angleson, J.K., MacMorris, M.A., Domke, E., Hutton, J.F., Schwartz, C., and Hutton, J.C. (2004). Dense core vesicle dynamics in Caenorhabditis elegans neurons and the role of kinesin UNC-104. Traffic 5, 544–559.

9. Lo, K.Y., Kuzmin, A., Unger, S.M., Petersen, J.D., and Silverman, M.A. (2011). KIF1A is the primary anterograde motor protein required for the axonal transport of dense-core vesicles in cultured hippocampal neurons. Neurosci Lett 491, 168–173.

10. Yonekawa, Y., Harada, A., Okada, Y., Funakoshi, T., Kanai, Y., Takei, Y., Terada, S., Noda, T., and Hirokawa, N. (1998). Defect in synaptic vesicle precursor transport and neuronal cell death in KIF1A motor protein-deficient mice. J Cell Biol 141, 431–441.

11. Pack-Chung, E., Kurshan, P.T., Dickman, D.K., and Schwarz, T.L. (2007). A Drosophila kinesin required for synaptic bouton formation and synaptic vesicle transport. Nat Neurosci 10, 980–989.

12. Gondre-Lewis, M.C., Park, J.J., and Loh, Y.P. (2012). Cellular mechanisms for the biogenesis and transport of synaptic and dense-core vesicles. Int Rev Cell Mol Biol 299, 27–115.

13. Takamori, S., Holt, M., Stenius, K., Lemke, E.A., Gronborg, M., Riedel, D., Urlaub, H., Schenck, S., Brugger, B., Ringler, P., et al. (2006). Molecular anatomy of a trafficking organelle. Cell 127, 831–846.

14. Li, P., Merrill, S.A., Jorgensen, E.M., and Shen, K. (2016). Two Clathrin Adaptor Protein Complexes Instruct Axon-Dendrite Polarity. Neuron 90, 564–580.

15. Ailion, M., Hannemann, M., Dalton, S., Pappas, A., Watanabe, S., Hegermann, J., Liu, Q., Han, H.F., Gu, M., Goulding, M.Q., et al. (2014). Two Rab2 interactors regulate dense-core vesicle maturation. Neuron 82, 167–180.

16. Sumakovic, M., Hegermann, J., Luo, L., Husson, S.J., Schwarze, K., Olendrowitz, C., Schoofs, L., Richmond, J., and Eimer, S. (2009). UNC-108/RAB-2 and its effector RIC-19 are involved in dense core vesicle maturation in Caenorhabditis elegans. J Cell Biol 186, 897–914.

17. Hummel, J.J.A., and Hoogenraad, C.C. (2021). Specific KIF1A-adaptor interactions control selective cargo recognition. J Cell Biol 220.

18. Emperador-Melero, J., Huson, V., van Weering, J., Bollmann, C., Fischer von Mollard, G., Toonen, R.F., and Verhage, M. (2018). Vti1a/b regulate synaptic vesicle and dense core vesicle secretion via protein sorting at the Golgi. Nat Commun 9, 3421.

19. De Camilli, P., and Jahn, R. (1990). Pathways to regulated exocytosis in neurons. Annu Rev Physiol 52, 625–645.

20. Burgess, J., Jauregui, M., Tan, J., Rollins, J., Lallet, S., Leventis, P.A., Boulianne, G.L., Chang, H.C., Le Borgne, R., Kramer, H., et al. (2011). AP-1 and clathrin are essential for secretory granule biogenesis in Drosophila. Mol Biol Cell 22, 2094–2105.

21. Akhmanova, A., and Steinmetz, M.O. (2008). Tracking the ends: a dynamic protein network controls the fate of microtubule tips. Nat Rev Mol Cell Biol 9, 309–322.

22. Hayashi, I., and Ikura, M. (2003). Crystal structure of the amino-terminal microtubule-binding domain of end-binding protein 1 (EB1). J Biol Chem 278, 36430–36434.

23. Honnappa, S., Gouveia, S.M., Weisbrich, A., Damberger, F.F., Bhavesh, N.S., Jawhari, H., Grigoriev, I., van Rijssel, F.J., Buey, R.M., Lawera, A., et al. (2009). An EB1-binding motif acts as a microtubule tip localization signal. Cell 138, 366–376.

24. Honnappa, S., John, C.M., Kostrewa, D., Winkler, F.K., and Steinmetz, M.O. (2005). Structural insights into the EB1-APC interaction. EMBO J 24, 261–269.

25. Maurer, S.P., Fourniol, F.J., Bohner, G., Moores, C.A., and Surrey, T. (2012). EBs recognize a nucleotide-dependent structural cap at growing microtubule ends. Cell 149, 371–382.

26. Leterrier, C., Vacher, H., Fache, M.P., d’Ortoli, S.A., Castets, F., Autillo-Touati, A., and Dargent, B. (2011). End-binding proteins EB3 and EB1 link microtubules to ankyrin G in the axon initial segment. Proc Natl Acad Sci U S A 108, 8826–8831.

27. Mattie, F.J., Stackpole, M.M., Stone, M.C., Clippard, J.R., Rudnick, D.A., Qiu, Y., Tao, J., Allender, D.L., Parmar, M., and Rolls, M.M. (2010). Directed microtubule growth, +TIPs, and kinesin-2 are required for uniform microtubule polarity in dendrites. Curr Biol 20, 2169–2177.

28. Yogev, S., Cooper, R., Fetter, R., Horowitz, M., and Shen, K. (2016). Microtubule Organization Determines Axonal Transport Dynamics. Neuron 92, 449–460.

29. Yang, C., Wu, J., de Heus, C., Grigoriev, I., Liv, N., Yao, Y., Smal, I., Meijering, E., Klumperman, J., Qi, R.Z., et al. (2017). EB1 and EB3 regulate microtubule minus end organization and Golgi morphology. J Cell Biol 216, 3179–3198.

30. Guedes-Dias, P., Nirschl, J.J., Abreu, N., Tokito, M.K., Janke, C., Magiera, M.M., and Holzbaur, E.L.F. (2019). Kinesin-3 Responds to Local Microtubule Dynamics to Target Synaptic Cargo Delivery to the Presynapse. Curr Biol 29, 268–282 e268.

31. Alves-Silva, J., Sanchez-Soriano, N., Beaven, R., Klein, M., Parkin, J., Millard, T.H., Bellen, H.J., Venken, K.J., Ballestrem, C., Kammerer, R.A., et al. (2012). Spectraplakins promote microtubule-mediated axonal growth by functioning as structural microtubule-associated proteins and EB1-dependent +TIPs (tip interacting proteins). J Neurosci 32, 9143–9158.

32. Moughamian, A.J., Osborn, G.E., Lazarus, J.E., Maday, S., and Holzbaur, E.L. (2013). Ordered recruitment of dynactin to the microtubule plus-end is required for efficient initiation of retrograde axonal transport. J Neurosci 33, 13190–13203.

33. Komarova, Y., De Groot, C.O., Grigoriev, I., Gouveia, S.M., Munteanu, E.L., Schober, J.M., Honnappa, S., Buey, R.M., Hoogenraad, C.C., Dogterom, M., et al. (2009). Mammalian end binding proteins control persistent microtubule growth. J Cell Biol 184, 691–706.

34. Schmidt, R., Fielmich, L.E., Grigoriev, I., Katrukha, E.A., Akhmanova, A., and van den Heuvel, S. (2017). Two populations of cytoplasmic dynein contribute to spindle positioning in C. elegans embryos. J Cell Biol 216, 2777–2793.

35. Klassen, M.P., and Shen, K. (2007). Wnt signaling positions neuromuscular connectivity by inhibiting synapse formation in C. elegans. Cell 130, 704–716.

36. White, J.G., Southgate, E., Thomson, J.N., and Brenner, S. (1986). The structure of the nervous system of the nematode Caenorhabditis elegans. Philos Trans R Soc Lond B Biol Sci 314, 1–340.

37. Hammarlund, M., Watanabe, S., Schuske, K., and Jorgensen, E.M. (2008). CAPS and syntaxin dock dense core vesicles to the plasma membrane in neurons. J Cell Biol 180, 483–491.

38. Morrison, L.M., Edwards, S.L., Manning, L., Stec, N., Richmond, J.E., and Miller, K.G. (2018). Sentryn and SAD Kinase Link the Guided Transport and Capture of Dense Core Vesicles in Caenorhabditis elegans. Genetics 210, 925–946.

39. Speese, S., Petrie, M., Schuske, K., Ailion, M., Ann, K., Iwasaki, K., Jorgensen, E.M., and Martin, T.F. (2007). UNC-31 (CAPS) is required for dense-core vesicle but not synaptic vesicle exocytosis in Caenorhabditis elegans. J Neurosci 27, 6150–6162.

40. van Keimpema, L., Kooistra, R., Toonen, R.F., and Verhage, M. (2017). CAPS-1 requires its C2, PH, MHD1 and DCV domains for dense core vesicle exocytosis in mammalian CNS neurons. Sci Rep 7, 10817.

41. Fares, H., and Greenwald, I. (2001). Genetic analysis of endocytosis in Caenorhabditis elegans: coelomocyte uptake defective mutants. Genetics 159, 133–145.

42. Brewer, J.C., Olson, A.C., Collins, K.M., and Koelle, M.R. (2019). Serotonin and neuropeptides are both released by the HSN command neuron to initiate Caenorhabditis elegans egg laying. PLoS Genet 15, e1007896.

43. Banerjee, N., Bhattacharya, R., Gorczyca, M., Collins, K.M., and Francis, M.M. (2017). Local neuropeptide signaling modulates serotonergic transmission to shape the temporal organization of C. elegans egg-laying behavior. PLoS Genet 13, e1006697.

44. Schwartz, M.L., and Jorgensen, E.M. (2016). SapTrap, a Toolkit for High-Throughput CRISPR/Cas9 Gene Modification in Caenorhabditis elegans. Genetics 202, 1277–1288.

45. Xuan, Z., Manning, L., Nelson, J., Richmond, J.E., Colon-Ramos, D.A., Shen, K., and Kurshan, P.T. (2017). Clarinet (CLA-1), a novel active zone protein required for synaptic vesicle clustering and release. Elife 6.

46. Akhmanova, A., Hoogenraad, C.C., Drabek, K., Stepanova, T., Dortland, B., Verkerk, T., Vermeulen, W., Burgering, B.M., De Zeeuw, C.I., Grosveld, F., et al. (2001). CLASPs Are CLIP-115 and −170 Associating Proteins Involved in the Regional Regulation of Microtubule Dynamics in Motile Fibroblasts. Cell 104, 923–935.

47. Tanenbaum, M.E., Macurek, L., van der Vaart, B., Galli, M., Akhmanova, A., and Medema, R.H. (2011). A complex of Kif18b and MCAK promotes microtubule depolymerization and is negatively regulated by Aurora kinases. Curr Biol 21, 1356–1365.

48. van der Vaart, B., Manatschal, C., Grigoriev, I., Olieric, V., Gouveia, S.M., Bjelic, S., Demmers, J., Vorobjev, I., Hoogenraad, C.C., Steinmetz, M.O., et al. (2011). SLAIN2 links microtubule plus end-tracking proteins and controls microtubule growth in interphase. J Cell Biol 193, 1083–1099.

49. Jiang, K., Toedt, G., Montenegro Gouveia, S., Davey, N.E., Hua, S., van der Vaart, B., Grigoriev, I., Larsen, J., Pedersen, L.B., Bezstarosti, K., et al. (2012). A Proteome-wide screen for mammalian SxIP motif-containing microtubule plus-end tracking proteins. Curr Biol 22, 1800–1807.

50. Luo, L., Hannemann, M., Koenig, S., Hegermann, J., Ailion, M., Cho, M.K., Sasidharan, N., Zweckstetter, M., Rensing, S.A., and Eimer, S. (2011). The Caenorhabditis elegans GARP complex contains the conserved Vps51 subunit and is required to maintain lysosomal morphology. Mol Biol Cell 22, 2564–2578.

51. Jaulin, F., and Kreitzer, G. (2010). KIF17 stabilizes microtubules and contributes to epithelial morphogenesis by acting at MT plus ends with EB1 and APC. J Cell Biol 190, 443–460.

52. Ligon, L.A., Shelly, S.S., Tokito, M., and Holzbaur, E.L. (2003). The microtubule plus-end proteins EB1 and dynactin have differential effects on microtubule polymerization. Mol Biol Cell 14, 1405–1417.

53. Kornakov, N., Mollers, B., and Westermann, S. (2020). The EB1-Kinesin-14 complex is required for efficient metaphase spindle assembly and kinetochore bi-orientation. J Cell Biol 219.

54. Morris, E.J., Nader, G.P., Ramalingam, N., Bartolini, F., and Gundersen, G.G. (2014). Kif4 interacts with EB1 and stabilizes microtubules downstream of Rho-mDia in migrating fibroblasts. PLoS One 9, e91568.

55. Stout, J.R., Yount, A.L., Powers, J.A., Leblanc, C., Ems-McClung, S.C., and Walczak, C.E. (2011). Kif18B interacts with EB1 and controls astral microtubule length during mitosis. Mol Biol Cell 22, 3070–3080.

56. Shin, H., Wyszynski, M., Huh, K.H., Valtschanoff, J.G., Lee, J.R., Ko, J., Streuli, M., Weinberg, R.J., Sheng, M., and Kim, E. (2003). Association of the kinesin motor KIF1A with the multimodular protein liprin-alpha. J Biol Chem 278, 11393–11401.

57. Wagner, O.I., Esposito, A., Kohler, B., Chen, C.W., Shen, C.P., Wu, G.H., Butkevich, E., Mandalapu, S., Wenzel, D., Wouters, F.S., et al. (2009). Synaptic scaffolding protein SYD-2 clusters and activates kinesin-3 UNC-104 in C. elegans. Proc Natl Acad Sci U S A 106, 19605–19610.

58. Gotz, T.W.B., Puchkov, D., Lysiuk, V., Lutzkendorf, J., Nikonenko, A.G., Quentin, C., Lehmann, M., Sigrist, S.J., and Petzoldt, A.G. (2021). Rab2 regulates presynaptic precursor vesicle biogenesis at the trans-Golgi. J Cell Biol 220.

59. Rizalar, F.S., Roosen, D.A., and Haucke, V. (2021). A Presynaptic Perspective on Transport and Assembly Mechanisms for Synapse Formation. Neuron 109, 27–41.

60. Zahavi, E.E., Hummel, J.J.A., Han, Y., Bar, C., Stucchi, R., Altelaar, M., and Hoogenraad, C.C. (2021). Combined kinesin-1 and kinesin-3 activity drives axonal trafficking of TrkB receptors in Rab6 carriers. Dev Cell 56, 494–508 e497.

61. Qu, X., Kumar, A., Blockus, H., Waites, C., and Bartolini, F. (2019). Activity-Dependent Nucleation of Dynamic Microtubules at Presynaptic Boutons Controls Neurotransmission. Curr Biol 29, 4231–4240 e4235.

62. Roth, D., Fitton, B.P., Chmel, N.P., Wasiluk, N., and Straube, A. (2018). Spatial positioning of EB family proteins at microtubule tips involves distinct nucleotide-dependent binding properties. J Cell Sci 132.

63. Lord, S.J., Velle, K.B., Mullins, R.D., and Fritz-Laylin, L.K. (2020). SuperPlots: Communicating reproducibility and variability in cell biology. J Cell Biol 219.

64. Brenner, S. (1974). The genetics of Caenorhabditis elegans. Genetics 77, 71–94.

65. Dokshin, G.A., Ghanta, K.S., Piscopo, K.M., and Mello, C.C. (2018). Robust Genome Editing with Short Single-Stranded and Long, Partially Single-Stranded DNA Donors in Caenorhabditis elegans. Genetics 210, 781–787.

66. Herbert, A.D., Carr, A.M., and Hoffmann, E. (2014). FindFoci: a focus detection algorithm with automated parameter training that closely matches human assignments, reduces human inconsistencies and increases speed of analysis. PLoS One 9, e114749.

